# High Pathogenicity Avian Influenza Virus (HPAIV) H5N1 clade 2.3.4.4b recovered from a kelp gull (*Larus dominicanus*) in the South Shetland Islands, Antarctica

**DOI:** 10.1101/2024.12.29.630510

**Authors:** Maria Ogrzewalska, Elisa Cavalcante Pereira, Ralph Eric Thijl Vanstreels, Emandi Campista, Leonardo Correa Junior, Larissa Macedo, Luciana Reis Appolinario, Martha Lima Brandão, Roberto Vilela, Wim Degrave, Fernando Couto Motta, Marilda Mendonca Siqueira, Paola Cristina Resende

**Author notes:** **Corresponding authors:** Paola Cristina Resende; Maria Ogrzewalska.

## Abstract

Whole-genome analysis of the earliest-detected High Pathogenicity Avian Influenza Virus (HPAIV) H5N1 clade 2.3.4.4b detected in Hannah Point, Antarctica (January 2024) reveals close relatedness to strains that circulated in pinnipeds and seabirds along the Atlantic coast of South America during the second half of 2023.

## Correspondence Letter

The global spread of the High Pathogenicity Avian Influenza Virus (HPAIV) subtype H5N1, specifically clade 2.3.4.4b, has resulted in extraordinary mortality levels among wild animals^1^. In 2022 and 2023, the virus led to widespread deaths among seabirds and marine mammals in South America^2,3^. In October 2023, HPAIV H5N1 was detected on the South Georgia subantarctic islands causing considerable mortalities in seabirds and marine mammals^4^. The first reports of wildlife mortality suspected to be due to HPAIV in the Antarctic region was reported in November 2023. Since then, several cases have been confirmed in the Antarctic Peninsula region^5^. No full genome sequences of HPAIV detected in the region have been made available to date. Here, we present the first full-genome data from the HPAIV H5N1 clade 2.3.4.4b from the Antarctic region, recovered from a kelp gull (*Larus dominicanus*) in the South Shetland Islands.

On January 8, 2024, cloacal swabs were collected from an adult kelp gull (*Larus dominicanus*) found deceased at Hannah Point on the southern coast of Livingston Island, South Shetland Islands, Antarctica (62°39′16″S 60°36′48″W), **Supplementary Figure 1**. The methods used to analyse these samples for HPAIV are described in **Supplementary Material**. Real time RT-PCR for Influenza targets revealed a cycle threshold (ct) of 24.17 for the target InfA (gene M), and of 36.60 and 35.97, respectively, for the targets H5a and H5b hemagglutinin gene. To our knowledge, this was the earliest HPAIV detection in Antarctica, with other detections in the Antarctic Peninsula and South Shetland Islands occurring in February and March 2024^6,7^. In turn, the arrival of these viruses in Antarctica was preceded by HPAIV detections on the southern tip of South America in June 2023^8^ and in the Falkland/Malvinas Islands and South Georgia in October 2023^4^.

Total viral RNA was recovered from the swab sample, and sequenced. The complete genome sequence was assembled and had 100% of coverage for all gene segments (**Supplementary Table 1**). Phylogenetic reconstruction of the hemagglutinin gene revealed that A/kelp gull/South Shetland Islands/10272/2024 is a HPAIV subtype H5N1 and belongs to the clade 2.3.4.4b of the Goose/Guangdong lineage (**Figure 1**), with the B3.2 genotype (*sensu* Youk et al. 2024^9^). The phylogenetic reconstruction of neuraminidase and internal genes are represented in **Supplementary Figure 2**. The genomes used for the phylogenetic reconstruction are listed in **Supplementary Table 2**.

**Figure 1.**
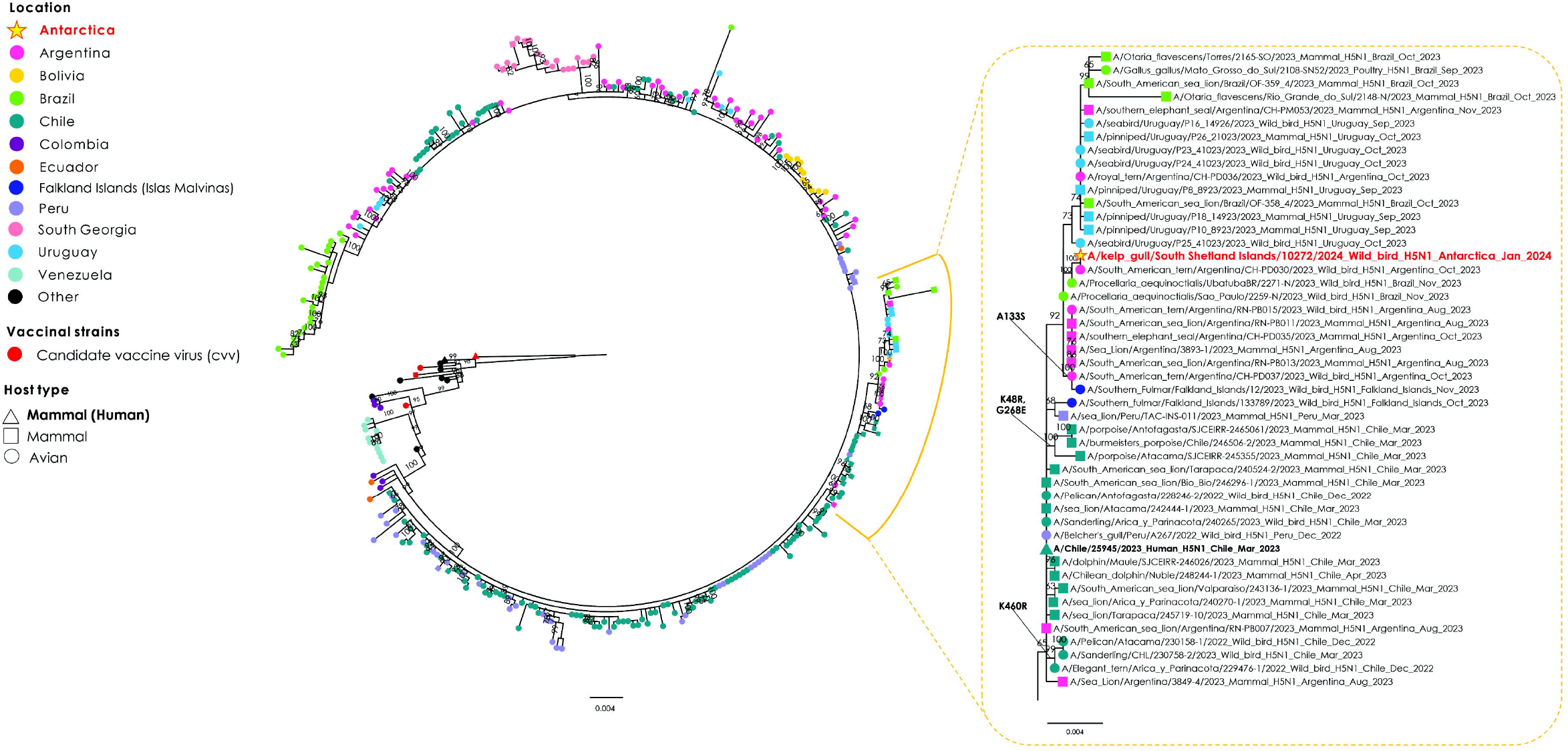
Phylogenetic tree based on maximum likelihood analysis of the hemagglutinin (HA) gene segment of Influenza A H5 clade 2.3.4.4b from a kelp gull identified in the South Shetland Islands, compared with HPAIV H5N1 sequences from South America and Antarctica, and selected strains from North America, Europe, and Africa. Node shape colours represent geographic region or country. Branch lengths indicate genetic divergence (scale bars provided). Bootstrap support values are shown next to each node.

The viral genome, along with their set of mutations, was evaluated at each of the M1, NA and PA sites relevant for antiviral susceptibility previously investigated, according to WHO guidelines. No amino acid changes associated with reduced neuraminidase inhibition were identified^10,11^. However, the mutation D701N in PB2, previously associated with enhanced polymerase activity in mammalian cells^12^, has not been investigated in phenotypic studies for this virus so far. Although only a single human case has been linked to this clade^13^, the possibility that these viruses harbour unidentified markers that may conferring an advantage in adapting to the human species cannot be ruled out. This underscores the importance of preserving collected samples to maintain viral integrity and enable virus isolation for functional and phenotypic testing.

Our phylogenetic analysis recovered the HPAIV strain detected in this study nested within a well-supported clade (bootstrap value: 100) comprising HPAIV strains found in birds and mammals across Colombia, Ecuador, Peru, Chile, Argentina, Uruguay, Brazil, Falkland/Malvinas Islands, and South Georgia since 2022. Within this group, genomic evidence shows that two different HPAIV H5N1 clades have emerged and spread in South America following the virus introduction to Peru: (i) the ‘avian’ clade was found in poultry and terrestrial and aquatic birds, with very few records in mammals, (ii) the ‘marine mammal’ clade was detected predominantly in aquatic mammals (sea lions, elephant seals, fur seals, dolphins, porpoises, and otters) with occasional spill-overs to seabirds^2^ and one human case^13^. HPAIV strains detected in seabirds (skuas, kelp gulls, and shags) and marine mammals (elephant seals and fur seals) in South Georgia belong to the ‘avian’ clade, whereas two HPAIV strains detected in seabirds (fulmars) in the Falkland/Malvinas Islands belonged to the ‘marine mammal’ clade^4^. The HPAIV strain recovered in this study nested within the ‘marine mammal’ clade (bootstrap value of 92), which thus far has been recorded in Peru, Chile, Argentina, Uruguay, Brazil, and the Falkland/Malvinas Islands. Conversely, the strain detected in this study does not appear to be a descendant nor closely related to the strains isolated from seabirds in the Falkland/Malvinas Islands, showing instead a higher similarity to the strains that circulated in pinnipeds and seabirds in Argentina, Uruguay, and Brazil. This suggests that the HPAIV was introduced from South America to Antarctica via an independent pathway, not mediated by South Georgia nor the Falkland/Malvinas Islands.

It is likely that seabirds (such as the kelp gull) or pinnipeds were responsible for the spread of HPAIV from South America to Antarctica. Kelp gulls are opportunistic predators and scavengers of birds and mammals and may engage in kleptoparasitism (i.e. deliberately taking the food from another animal) of other seabirds^14^. The species is widely distributed in the Southern Hemisphere, including South America and the Antarctic Peninsula, and during the austral winter the populations breeding in the Antarctic Peninsula often disperse to the north, reaching the southern tip of Argentina and Chile^14^. Considering that kelp gulls were frequently seen scavenging pinniped carcasses during HPAIV outbreaks in South America^2^ and are known to interact with other seabirds through predation or kleptoparasitism^14^, it is unsurprising that there were several HPAIV detections in kelp gulls in Peru, Chile, and Argentina in 2023^8^. It is therefore plausible that kelp gulls that breed in the Antarctic Peninsula exposed to HPAIV while wintering in South America may have carried the virus when returning to Antarctica. Alternatively, it is also possible that pinnipeds, particularly southern elephant seals (*Mirounga leonina*), played a direct role in the spread of the virus to Antarctica. Genetic studies suggest significant connectivity between the elephant seal populations in Argentina and the South Shetland Islands^15^. Considering that the species was heavily impacted by HPAIV (*c*. 95% pup mortality) in Argentina in October and November 2023^2^, it is conceivable that an elephant seal exposed to HPAIV in Argentina may have carried the virus to Antarctica and then locally transmitted it to other wildlife species (such as the kelp gull in this report) after coming ashore. This would be consistent with the fact that the HPAIV detection occurred at Hannah Point in January, which is when this site becomes a regional hotspot for moulting southern elephant seals^16^. Additional data on the genomes of other HPAIV strains from Antarctic wildlife are needed to provide more robust inferences about the pathways of introduction and spread of HPAIV to the Antarctica continent.

The detection of HPAIV H5N1 in Antarctica is an acute concern from an environmental perspective. It is estimated that more than 500,000 seabirds and 50,000 marine mammals died from HPAIV infection in South America^17^. If similar impacts emerge in Antarctica, the consequences for coastal and marine Antarctic ecosystems could be disastrous. Continued research, monitoring the spread and the impacts of these viruses in Antarctica in the coming years, will therefore be of paramount importance. Additionally, implementation of strict biosafety protocols and control measures will be crucial to prevent human-mediated spread of HPAIV H5N1 in the region and to mitigate zoonotic disease risks. Considering that transmission is bidirectional and that genetic reassortment between HPAIV and human influenza viruses is possible, it would also be highly recommended that people approaching or working with Antarctic wildlife should be vaccinated against seasonal influenza and, if possible, against influenza A (H5N1).

## ACKNOWLEDGMENTS

We thank the FIO-ANTAR Working Group, especially Sandra Soares and Lucia Marques; the Brazilian Antarctic Program – PROANTAR and the Vice-Presidency of Production and Innovation in Health (Fiocruz VPPIS), as well as the Norwegian Polar Institute’s Quantarctica package.

We also acknowledge the submitters of Global Initiative on Sharing All Influenza Data (GISAID) for the EpiFlu database, and other sequence databases which were used to share gene sequences and associated information (**Supplementary Table 3**).

We also like to thank the financial support of the PROGRAMA ANTÁRTICO BRASILEIRO – PROANTAR, Conselho Nacional de Desenvolvimento Científico e Tecnológico (CNPq)/MCTIC/CAPES/FNDCT Nº 21/2018 – PROANTAR (442646/2018-6), Reference laboratories from the Oswaldo Cruz Foundation (CVSLR/FIOCRUZ), and the Carlos Chagas Filho Foundation for Supporting Research in the State of Rio de Janeiro FAPERJ Nº E-26/010/002546/2019; CNPq CABBIO (Grant number 423857/2021-5, PCR) and the CNPq productivity research fellowship (311759/2022-0, PCR). Elisa Pereira is a postdoc supported by INOVA/IOC/FIOCRUZ.

## AUTHOR APPROVALS

All authors have seen and approved the manuscript, and it hasn’t been accepted or published elsewhere.

