## Supplementary material for "High Pathogenicity Avian Influenza Virus (HPAIV) H5N1 clade 2.3.4.4b recovered from a kelp gull (*Larus dominicanus*) in the South Shetland Islands, Antarctica"

### Methods

Sampling was performed as part of the FIOANTAR project conducted by the Oswaldo Cruz Foundation (FIOCRUZ) as part of the Brazilian Antarctic Program (PROANTAR), with logistical support from the Brazilian Navy and in adherence to strict biosafety protocols. The sample was stored at –80ºC during transport to the Laboratory of Respiratory Viruses, Exanthematous Viruses, Enteroviruses, and Viral Emergencies (LVRE) at Oswaldo Cruz Institute (IOC), where RNA was extracted using the QIAamp Viral RNA mini kit (QIAgen). Real-time RT-PCR detection and subtyping was performed for different influenza A virus targets InfA from gene M and H5a and H5b from hemagglutinin gene. Whole-genome sequencing was recovered by the Multisegmented Reverse Transcription-PCR (M-RT-PCR) protocol (Zhou et al., 2009) and assembled by Iterative Refinement Meta-Assembler (IRMA) (Shepard et al., 2016) . The whole genome recovered A/kelp_gull/South Shetland Islands/10272/2024 was uploaded to the EpiFlu database from GISAID (accession number EPI_ISL_19589149). The genome metrics are provided in **Supplementary Table 1**. The Basic Local Alignment Search Tool (BLAST) was used to identify similar genomes. For each genomic segment, a BLAST search retrieved the top 100 sequences. Additionally, all sequences from clade 2.4.4.4b of influenza H5N1 in South America, available on GISAID and GenBank, were downloaded. After removing duplicates and redundant entries, the datasets were curated to retain only representative sequences based on host, geographic origin, and collection period. Four HA and NA sequences from candidate vaccine viruses (CVVs) were also included in the analysis: A/American Wigeon/South Carolina/22-000345-001/2021 (EPI_ISL_18133029); A/Fujian Sanyuan/21099/2017 (EPI_ISL_8767096); A/chicken/Ghana/AVL-763_21VIR7050-39/2021 (EPI_ISL_16997921); and A/Ezo Red Fox/Hokkaido/1/2022 (EPI_ISL_12174842) (https://www.who.int/teams/global-influenza-programme/vaccines/who-recommendations/zoonotic-influenza-viruses-and-candidate-vaccine-viruses).

Multiple sequence alignment was performed using MAFFT v7.475 (Katoh & Standley, 2013) , followed by Maximum Likelihood (ML) phylogenetic analysis with IQ-TREE2 v2.3.6 (Nguyen et al., 2015) . The best-fitting nucleotide substitution model was selected using the ModelFinder tool within IQ-TREE2. Branch support was assessed by the ultra-fast bootstrap with 5,000 replicates (Minh et al., 2013) . Phylogenetic trees were visualized and annotated using FigTree v1.4.4 (http://tree.bio.ed.ac.uk/software/figtree/) and further refined with CorelDRAW Graphics Suite (Corel Corporation). Amino acid comparisons and identity analyses were conducted using FluSurver (https://flusurver.bii.a-star.edu.sg/), while amino acid branch annotations were performed with TreeSub (https://github.com/tamuri/treesub).

**Supplementary Table 1.** Sequencing metrics of the High Pathogenicity Avian Influenza Virus (HPAIV) H5N1 clade 2.3.4.4b whole genome recovered from a deceased *Larus dominicanus* in the South Shetland Islands, Antarctic.

| Genomic features | HA | NA | PB2 | PB1 | PA | NP | MP | NS |
|---|---|---|---|---|---|---|---|---|
| Number of reads | 61,420 | 24,123 | 10,148 | 10,903 | 23,897 | 32,391 | 84,128 | 36,547 |
| Coverage | 100% (5,418x) | 100% (2,584) | 100% (692x) | 100% (779x) | 100% (1,667x) | 100% (3,239x) | 100% (12,619x) | 100% (6,840x) |

**Supplementary Figure 1.** Geographic location of Hannah Point, on the southern coast of Livingston Island in the South Shetland Islands, Antarctica (62°39′16″S, 60°36′48″W), where a deceased kelp gull (*Larus dominicanus*) was found infected with the high pathogenicity avian influenza virus (HPAIV) subtype H5N1. The map was created using ArcGIS Pro (ESRI).

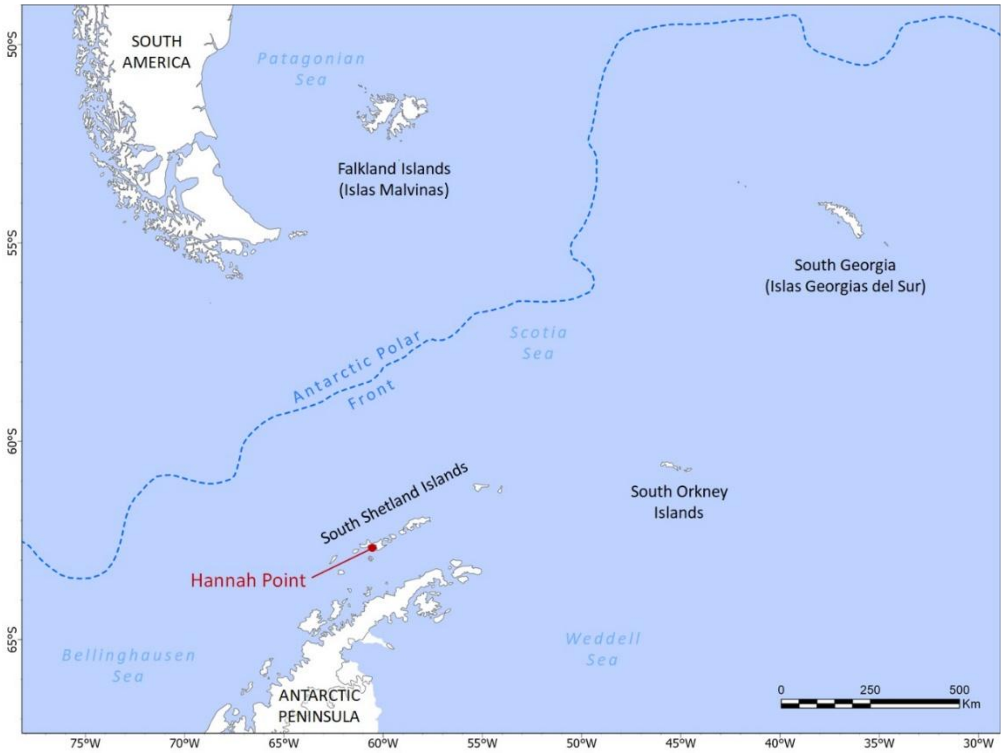

**Supplementary Figure 2.** Phylogenetic tree based on maximum likelihood analysis of the PB2 **(A)** and NA **(B)** gene segments from a kelp gull identified in the South Shetland Islands, compared with HPAI H5N1 sequences from South America and Antarctica, and selected strains from North America, Europe, and Africa used as outgroups. Node shape colours represent geographic region or country. Branch lengths indicate genetic divergence (scale bars provided). Bootstrap support values are shown next to each node.

**A**

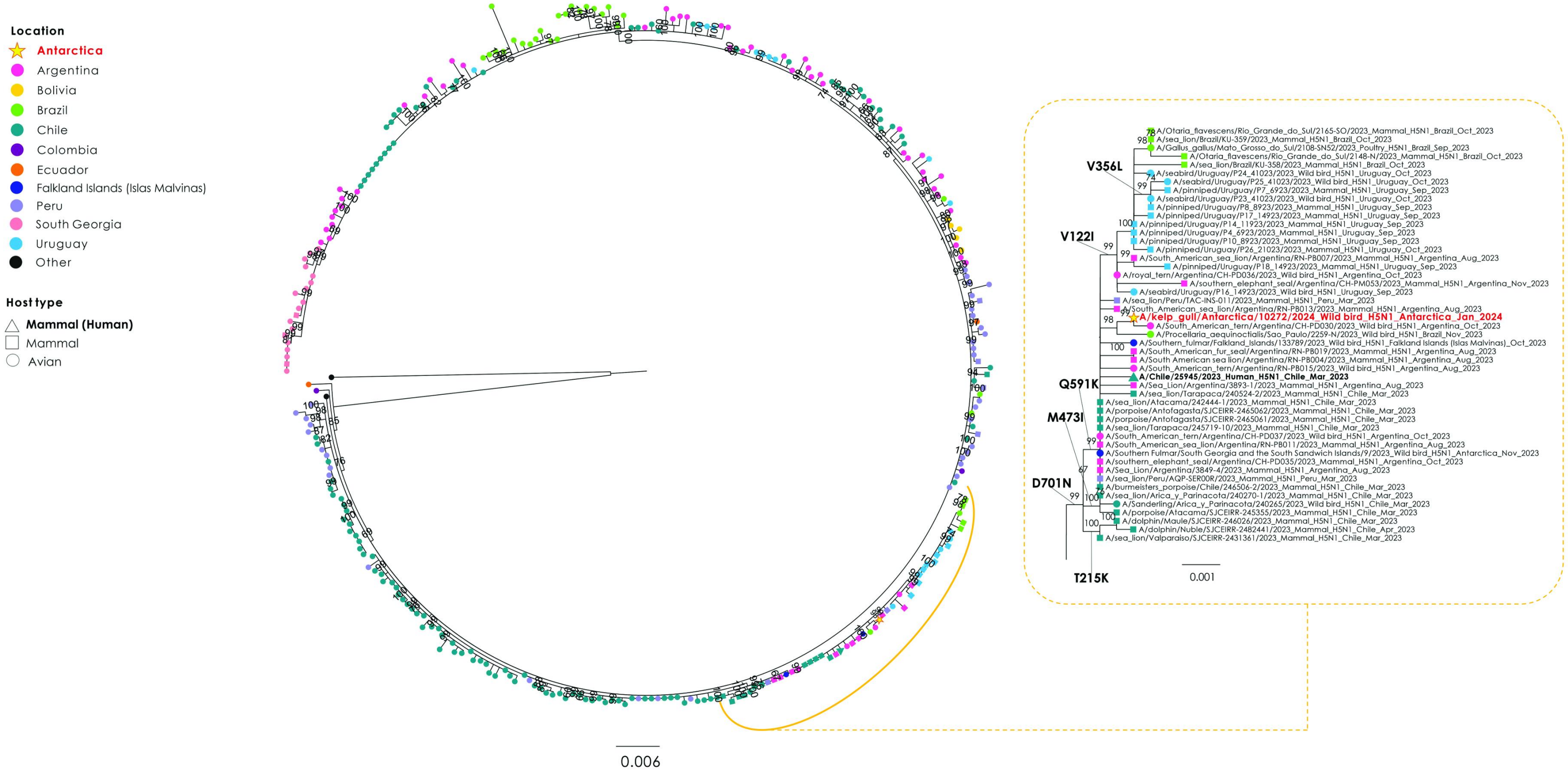

**B**

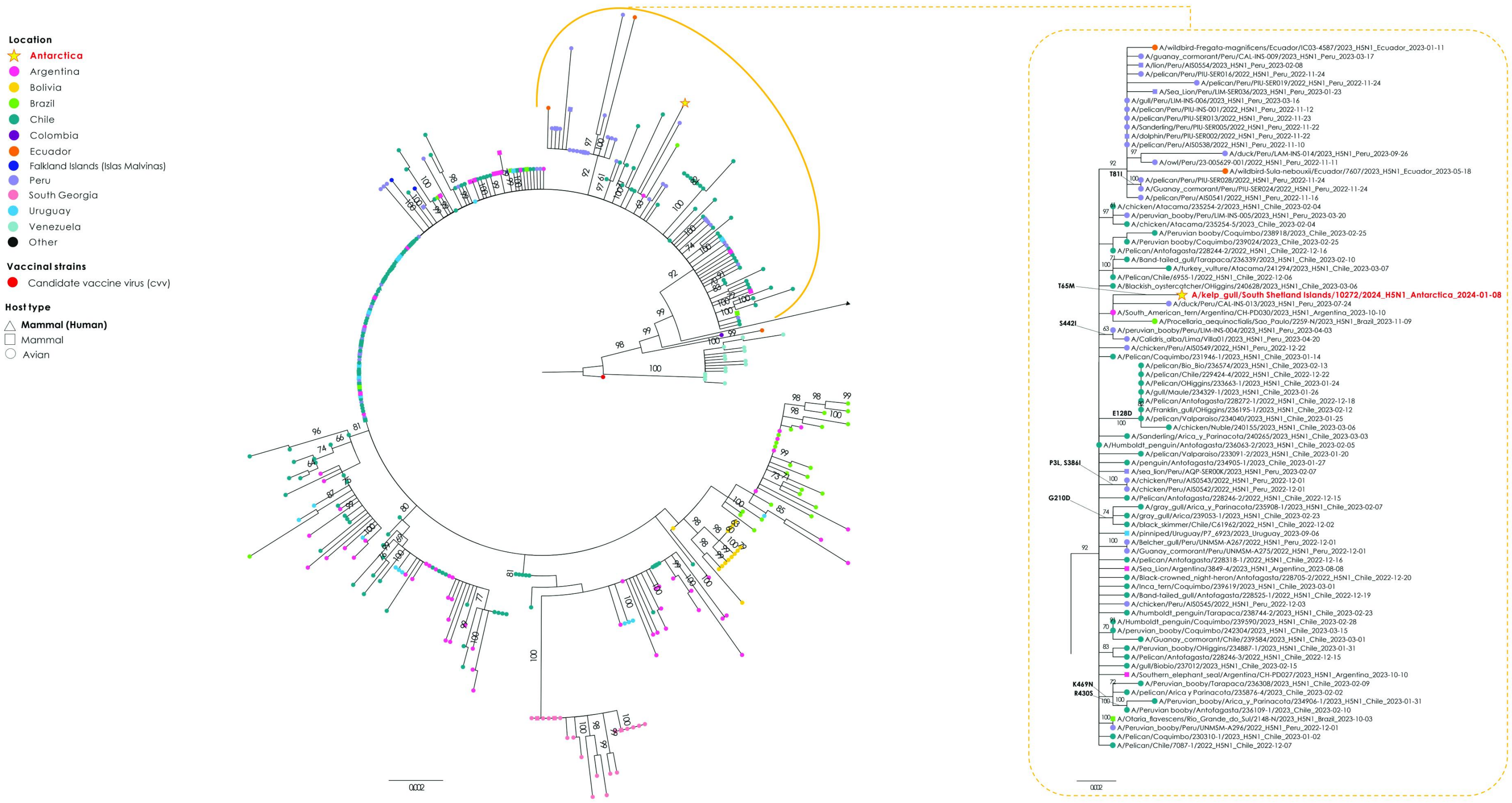
Location
Antarctica
Argentina
Bolivia
Brazil
Chile
Colombia
Ecuador
Falkland Islands (Islas Malvinas)
Peru
South Georgia
Uruguay
Venezuela
Other
Vaccinal strains
Candidate vaccine virus (cvv)
Host type
Mammal (Human)
Mammal
Avian
A/wildbird-Fregata-magnificens/Ecuador/IC03-4587/2023_H5N1_Ecuador_2023-01-11
A/guanay_cormorant/Peru/CAL-INS-009/2023_H5N1_Peru_2023-03-17
A/lion/Peru/AIS0554/2023_H5N1_Peru_2023-02-08
A/pelican/Peru/PIU-SER016/2022_H5N1_Peru_2022-11-24
A/pelican/Peru/PIU-SER019/2022_H5N1_Peru_2022-11-24
A/Sea_Lion/Peru/LIM-SER036/2023_H5N1_Peru_2023-01-23
A/gull/Peru/LIM-INS-006/2023_H5N1_Peru_2023-03-16
A/pelican/Peru/PIU-INS-001/2022_H5N1_Peru_2022-11-12
A/pelican/Peru/PIU-SER013/2022_H5N1_Peru_2022-11-23
A/Sanderling/Peru/PIU-SER005/2022_H5N1_Peru_2022-11-22
A/dolphin/Peru/PIU-SER002/2022_H5N1_Peru_2022-11-22
A/pelican/Peru/AIS0538/2022_H5N1_Peru_2022-11-10
A/duck/Peru/LAM-INS-014/2023_H5N1_Peru_2023-09-26
A/owl/Peru/23-005629-001/2022_H5N1_Peru_2022-11-11
A/wildbird-Sula-nebouxii/Ecuador/7607/2023_H5N1_Ecuador_2023-05-18
T81I
A/pelican/Peru/PIU-SER028/2022_H5N1_Peru_2022-11-24
A/Guanay_cormorant/Peru/PIU-SER024/2022_H5N1_Peru_2022-11-24
A/pelican/Peru/AIS0541/2022_H5N1_Peru_2022-11-16
A/chicken/Atacama/235254-2/2023_H5N1_Chile_2023-02-04
A/peruvian_booby/Peru/LIM-INS-005/2023_H5N1_Peru_2023-03-20
A/chicken/Atacama/235254-5/2023_Chile_2023-02-04
A/Peruvian booby/Coquimbo/238918/2023_Chile_2023-02-25
A/Peruvian booby/Coquimbo/239024/2023_Chile_2023-02-25
A/Pelican/Antofagasta/228244-2/2022_H5N1_Chile_2022-12-16
A/Band-tailed_gull/Tarapaca/236339/2023_H5N1_Chile_2023-02-10
A/turkey_vulture/Atacama/241294/2023_H5N1_Chile_2023-03-07
A/Pelican/Chile/6955-1/2022_H5N1_Chile_2022-12-06
A/Blackish_oystercatcher/OHiggins/240628/2023_H5N1_Chile_2023-03-06
T65M
A/kelp_gull/South Shetland Islands/10272/2024_H5N1_Antarctica_2024-01-08
A/duck/Peru/CAL-INS-013/2023_H5N1_Peru_2023-07-24
A/South_American_tern/Argentina/CH-PD030/2023_H5N1_Argentina_2023-10-10
A/Procellaria_aequinoctialis/Sao_Paulo/2259-N/2023_H5N1_Brazil_2023-11-09
S442I
A/peruvian_booby/Peru/LIM-INS-004/2023_H5N1_Peru_2023-04-03
A/Calidris_alba/Lima/Villa01/2023_H5N1_Peru_2023-04-20
A/chicken/Peru/AIS0549/2022_H5N1_Peru_2022-12-22
A/Pelican/Coquimbo/231946-1/2023_H5N1_Chile_2023-01-14
A/pelican/Bio_Bio/236574/2023_H5N1_Chile_2023-02-13
A/pelican/Chile/229424-4/2022_H5N1_Chile_2022-12-22
A/Pelican/OHiggins/233663-1/2023_H5N1_Chile_2023-01-24
A/gull/Maule/234329-1/2023_H5N1_Chile_2023-01-26
A/Pelican/Antofagasta/228272-1/2022_H5N1_Chile_2022-12-18
A/Franklin_gull/OHiggins/236195-1/2023_H5N1_Chile_2023-02-12
E128D
A/pelican/Valparaiso/234040/2023_H5N1_Chile_2023-01-25
A/chicken/Nuble/240155/2023_H5N1_Chile_2023-03-06
A/Sanderling/Arica_y_Parinacota/240265/2023_H5N1_Chile_2023-03-03
A/Humboldt_penguin/Antofagasta/236063-2/2023_H5N1_Chile_2023-02-05
A/pelican/Valparaiso/233091-2/2023_H5N1_Chile_2023-01-20
P3L, S386I
A/penguin/Antofagasta/234905-1/2023_Chile_2023-01-27
A/sea_lion/Peru/AQP-SER00K/2023_H5N1_Peru_2023-02-07
A/chicken/Peru/AIS0543/2022_H5N1_Peru_2022-12-01
A/chicken/Peru/AIS0542/2022_H5N1_Peru_2022-12-01
G210D
A/Pelican/Antofagasta/228246-2/2022_H5N1_Chile_2022-12-15
A/gray_gull/Arica_y_Parinacota/235908-1/2023_H5N1_Chile_2023-02-07
A/gray_gull/Arica/239053-1/2023_H5N1_Chile_2023-02-23
A/black_skimmer/Chile/C61962/2022_H5N1_Chile_2022-12-02
A/pinniped/Uruguay/P7_6923/2023_Uruguay_2023-09-06
A/Belcher_gull/Peru/UNMSM-A267/2022_H5N1_Peru_2022-12-01
A/Guanay_cormorant/Peru/UNMSM-A275/2022_H5N1_Peru_2022-12-01
A/pelican/Antofagasta/228318-1/2022_H5N1_Chile_2022-12-16
A/Sea_Lion/Argentina/3849-4/2023_H5N1_Argentina_2023-08-08
A/Black-crowned_night-heron/Antofagasta/228705-2/2022_H5N1_Chile_2022-12-20
A/Inca_tern/Coquimbo/239619/2023_H5N1_Chile_2023-03-01
A/Band-tailed_gull/Antofagasta/228525-1/2022_H5N1_Chile_2022-12-19
A/chicken/Peru/AIS0545/2022_H5N1_Peru_2022-12-03
A/humboldt_penguin/Tarapaca/238744-2/2023_H5N1_Chile_2023-02-23
A/Humboldt_penguin/Coquimbo/239590/2023_H5N1_Chile_2023-02-28
A/peruvian_booby/Coquimbo/242304/2023_H5N1_Chile_2023-03-15
A/Guanay_cormorant/Chile/239584/2023_H5N1_Chile_2023-03-01
A/Peruvian_booby/OHiggins/234887-1/2023_H5N1_Chile_2023-01-31
A/Pelican/Antofagasta/228246-3/2022_H5N1_Chile_2022-12-15
A/gull/Biobio/237012/2023_H5N1_Chile_2023-02-15
A/Southern_elephant_seal/Argentina/CH-PD027/2023_H5N1_Argentina_2023-10-10
K469N
R430S
A/Peruvian_booby/Tarapaca/236308/2023_H5N1_Chile_2023-02-09
A/pelican/Arica y Parinacota/235876-4/2023_Chile_2023-02-02
A/Peruvian_booby/Arica_y_Parinacota/234906-1/2023_H5N1_Chile_2023-01-31
A/Peruvian booby/Antofagasta/236109-1/2023_Chile_2023-02-10
A/Otaria_flavescens/Rio_Grande_do_Sul/2148-N/2023_H5N1_Brazil_2023-10-03
A/Peruvian_booby/Peru/UNMSM-A296/2022_H5N1_Peru_2022-12-01
A/Pelican/Coquimbo/230310-1/2023_H5N1_Chile_2023-01-02
A/Pelican/Chile/7087-1/2022_H5N1_Chile_2022-12-07

**Supplementary Table 2.** Accession codes for gene sequences used in phylogenetic analyses and mutational profiling, obtained from the Global Initiative on Sharing All Influenza Data (GISAID) and GenBank NCBI.

| Isolate Name | Accession Code |
|---|---|
| A/kelp_gull/South Shetland Islands/10272/2024 | EPI_ISL_19589149 |
| A/Astrakhan/3212/2020 | EPI_ISL_1038924 |
| A/Ezo red fox/Hokkaido/1/2022 | EPI_ISL_12174842 |
| A/chicken/Wyoming/22-009599-002/2022 | EPI_ISL_13009694 |
| A/Pelican/Venezuela/Pel4/2022 | EPI_ISL_16013753 |
| A/chicken/Ecuador/02/2022 | EPI_ISL_16157545 |
| A/pelican/Peru/PIU-INS-001/2022 | EPI_ISL_16249274 |
| A/chicken/Peru/LAM-INS-002/2022 | EPI_ISL_16249681 |
| A/chicken/Peru/LIM-INS-003/2022 | EPI_ISL_16249730 |
| A/Pelican/Venezuela/Inf9/2022 | EPI_ISL_16631973 |
| A/gray gull/Chile/C61947/2022 | EPI_ISL_16891400 |
| A/Peruvian pelican/Chile/C61740/2022 | EPI_ISL_16891401 |
| A/black skimmer/Chile/C61962/2022 | EPI_ISL_16891402 |
| A/chicken/Ghana/AVL-763_21VIR7050-39/2021 | EPI_ISL_16997921 |
| A/Pelican/Chile/6955-1/2022 | EPI_ISL_17011956 |
| A/Pelican/Chile/6955-3/2022 | EPI_ISL_17011957 |
| A/Gull/Chile/227023-3/2022 | EPI_ISL_17011958 |
| A/Pelican/Chile/6958-1/2022 | EPI_ISL_17011959 |
| A/Pelican/Chile/6618-1/2022 | EPI_ISL_17011962 |
| A/Pelican/Chile/6924-1/2022 | EPI_ISL_17011963 |
| A/Gull/Chile/227023-2/2022 | EPI_ISL_17011964 |
| A/Pelican/Chile/7087-1/2022 | EPI_ISL_17012018 |
| A/Pelican/Chile/227023-1/2022 | EPI_ISL_17050893 |
| A/chicken/Bolivar/ICA-3500/2022 | EPI_ISL_17353507 |
| A/chicken/Choco/ICA-3502/2022 | EPI_ISL_17353508 |
| A/chicken/Choco/ICA-3504/2022 | EPI_ISL_17353509 |
| A/chicken/Cordoba/ICA-3499/2022 | EPI_ISL_17353510 |
| A/chicken/Magdalena/ICA-3503/2022 | EPI_ISL_17353838 |
| A/duck/Choco/ICA-3501/2022 | EPI_ISL_17353839 |
| A/Chile/25945/2023 | EPI_ISL_17468386 |
| A/pelican/Peru/A074/2022 | EPI_ISL_17477219 |
| A/gull/Peru/A102/2022 | EPI_ISL_17477220 |
| A/falcon/Peru/A273/2022 | EPI_ISL_17477223 |
| A/chicken/Peru/23-005607-001/2022 | EPI_ISL_17660070 |
| A/chicken/Peru/23-005607-002/2022 | EPI_ISL_17660071 |
| A/chicken/Peru/23-005607-003/2022 | EPI_ISL_17660072 |

| | |
|---|---|
| A/owl/Peru/23-005629-001/2022 | EPI_ISL_17660073 |
| A/brown booby/Peru/23-005629-003/2022 | EPI_ISL_17660074 |
| A/peruvian_booby/Peru/LIM-INS-004/2023 | EPI_ISL_17777525 |
| A/peruvian_booby/Peru/LIM-INS-005/2023 | EPI_ISL_17777526 |
| A/gull/Peru/LIM-INS-006/2023 | EPI_ISL_17777527 |
| A/peruvian_booby/Peru/CAL-INS-007/2023 | EPI_ISL_17777528 |
| A/peruvian_booby/Peru/CAL-INS-008/2023 | EPI_ISL_17777529 |
| A/guanay_cormorant/Peru/CAL-INS-009/2023 | EPI_ISL_17777530 |
| A/sea_lion/Peru/TAC-INS-010/2023 | EPI_ISL_17777531 |
| A/sea_lion/Peru/TAC-INS-011/2023 | EPI_ISL_17777532 |
| A/peruvian_booby/Peru/LIM-INS-012/2023 | EPI_ISL_17777533 |
| A/chicken/Peru/AIS0546/2022 | EPI_ISL_17805986 |
| A/chicken/Peru/AIS0539/2022 | EPI_ISL_17805987 |
| A/chicken/Peru/AIS0540/2022 | EPI_ISL_17805988 |
| A/chicken/Peru/AIS0542/2022 | EPI_ISL_17805989 |
| A/chicken/Peru/AIS0543/2022 | EPI_ISL_17805990 |
| A/chicken/Peru/AIS0544/2022 | EPI_ISL_17805991 |
| A/chicken/Peru/AIS0545/2022 | EPI_ISL_17805992 |
| A/chicken/Peru/AIS0547/2022 | EPI_ISL_17805993 |
| A/chicken/Peru/AIS0548/2022 | EPI_ISL_17805994 |
| A/chicken/Peru/AIS0550/2022 | EPI_ISL_17805995 |
| A/chicken/Peru/AIS0549/2022 | EPI_ISL_17805996 |
| A/avian/Peru/AISA0446/2022 | EPI_ISL_17805997 |
| A/avian/Peru/AISA0451/2022 | EPI_ISL_17805998 |
| A/lion/Peru/AIS0554/2023 | EPI_ISL_17805999 |
| A/pelican/Peru/AIS0538/2022 | EPI_ISL_17806000 |
| A/chicken/Peru/AIS0551/2022 | EPI_ISL_17806001 |
| A/pelican/Peru/AIS0541/2022 | EPI_ISL_17806002 |
| A/pelican/Peru/AISA0464/2022 | EPI_ISL_17806003 |
| A/Black Skimmer/Maule/240379/2023 | EPI_ISL_17885849 |
| A/goose/Araucania/239189-1/2023 | EPI_ISL_17885869 |
| A/chicken/OHiggins/241252-6/2023 | EPI_ISL_17885870 |
| A/chicken/OHiggins/241252-3/2023 | EPI_ISL_17885871 |
| A/chicken/OHiggins/241252-1/2023 | EPI_ISL_17885872 |
| A/chicken/Nuble/241681-1/2023 | EPI_ISL_17885873 |
| A/chicken/Nuble/241557-1/2023 | EPI_ISL_17885874 |
| A/chicken/Nuble/240684/2023 | EPI_ISL_17885876 |
| A/chicken/Nuble/240155/2023 | EPI_ISL_17885877 |
| A/chicken/Nuble/239136/2023 | EPI_ISL_17885878 |
| A/Pelican/OHiggins/233721-1/2023 | EPI_ISL_17885912 |
| A/Pelican/OHiggins/233663-1/2023 | EPI_ISL_17885915 |

| | |
|---|---|
| A/Pelican/Nuble/233947-2/2023 | EPI_ISL_17885918 |
| A/Pelican/Maule/231155-2/2023 | EPI_ISL_17885921 |
| A/Pelican/Coquimbo/231946-1/2023 | EPI_ISL_17885922 |
| A/Pelican/Coquimbo/230310-1/2023 | EPI_ISL_17885923 |
| A/Pelican/Atacama/230158-2/2022 | EPI_ISL_17885924 |
| A/Pelican/Atacama/230158-1/2022 | EPI_ISL_17885925 |
| A/Pelican/Atacama/229450-2/2022 | EPI_ISL_17885926 |
| A/Pelican/Atacama/229424-2/2022 | EPI_ISL_17885927 |
| A/pelican/Antofagasta/228318-1/2022 | EPI_ISL_17885928 |
| A/Pelican/Antofagasta/228272-1/2022 | EPI_ISL_17885929 |
| A/pelican/Valparaiso/233091-1/2023 | EPI_ISL_17885930 |
| A/Pelican/Antofagasta/228246-3/2022 | EPI_ISL_17885931 |
| A/pelican/Valparaiso/233418-1/2023 | EPI_ISL_17885932 |
| A/Pelican/Antofagasta/228246-2/2022 | EPI_ISL_17885934 |
| A/Pelican/Tarapaca/227436-2/2022 | EPI_ISL_17885942 |
| A/Pelican/Antofagasta/228244-2/2022 | EPI_ISL_17885944 |
| A/chicken/Araucania/241914-1/2023 | EPI_ISL_17885945 |
| A/chicken/Araucania/241892-2/2023 | EPI_ISL_17885946 |
| A/humboldt penguin/Tarapaca/238744-2/2023 | EPI_ISL_17885947 |
| A/Humboldt penguin/Coquimbo/239590/2023 | EPI_ISL_17885948 |
| A/chicken/Araucania/240481-1/2023 | EPI_ISL_17885949 |
| A/Great egret/Araucania/240518/2023 | EPI_ISL_17885950 |
| A/Gray gull/Tarapaca/232825-1/2023 | EPI_ISL_17885951 |
| A/chicken/Araucania/239569-2/2023 | EPI_ISL_17885952 |
| A/Elegant tern/Tarapaca/229133-1/2022 | EPI_ISL_17885953 |
| A/chicken/Araucania/239569-1/2023 | EPI_ISL_17885954 |
| A/Elegant tern/Arica y Parinacota/229476-1/2022 | EPI_ISL_17885955 |
| A/pelican/Valparaiso/233091-2/2023 | EPI_ISL_17885956 |
| A/duck/Maule/240466-1/2023 | EPI_ISL_17885957 |
| A/chicken/Araucania/239189-2/2023 | EPI_ISL_17885958 |
| A/duck/Araucania/241914-2/2023 | EPI_ISL_17885959 |
| A/whimbrel/Coquimbo/239964/2023 | EPI_ISL_17885960 |
| A/duck/Araucania/240481-2/2023 | EPI_ISL_17885961 |
| A/turkey/Nuble/241568-1/2023 | EPI_ISL_17885962 |
| A/duck/Araucania/239189-3/2023 | EPI_ISL_17885963 |
| A/turkey/Nuble/240489-1/2023 | EPI_ISL_17885964 |
| A/Chiloe wigeon/OHiggins/240893-2/2023 | EPI_ISL_17885965 |
| A/turkey/Araucania/241892-3/2023 | EPI_ISL_17885966 |
| A/Blackish oystercatcher/OHiggins/240628/2023 | EPI_ISL_17885967 |
| A/Turkey vulture/Valparaiso/230187-1/2022 | EPI_ISL_17885968 |
| A/heron/Antofagasta/228705-3/2022 | EPI_ISL_17885969 |

| | |
|---|---|
| A/Turkey vulture/Antofagasta/228252-1/2022 | EPI_ISL_17885970 |
| A/heron/Antofagasta/228705-2/2022 | EPI_ISL_17885971 |
| A/Kelp gull/Maule/239349/2023 | EPI_ISL_17885972 |
| A/South American tern/Maule/238507/2023 | EPI_ISL_17885973 |
| A/sea lion/Tarapaca/240524-2/2023 | EPI_ISL_17885975 |
| A/sea lion/Arica y Parinacota/240270-1/2023 | EPI_ISL_17885976 |
| A/Sanderling/Arica y Parinacota/240265/2023 | EPI_ISL_17885978 |
| A/Sanderling/Arica y Parinacota/230758-1/2022 | EPI_ISL_17885980 |
| A/pelican/Valparaiso/234040/2023 | EPI_ISL_17885982 |
| A/pelican/Valparaiso/233450-1/2023 | EPI_ISL_17885983 |
| A/pelican/Valparaiso/233447-2/2023 | EPI_ISL_17885985 |
| A/whimbrel/Valparaiso/239946/2023 | EPI_ISL_17885987 |
| A/wildbird-Fregata-magnificens/Ecuador/IC03-4587/2023 | EPI_ISL_17973443 |
| A/wildbird-Fregata-magnificens/Ecuador/IC06-4590/2023 | EPI_ISL_17973458 |
| A/Black-crowned night-heron/Antofagasta/228705-2/2022 | EPI_ISL_18005786 |
| A/guanay cormorant/Peru/PIU-024/2022 | EPI_ISL_18054500 |
| A/sanderling/Peru/PIU-005/2022 | EPI_ISL_18054501 |
| A/Sea Lion/Peru/LIM-SER036/2023 | EPI_ISL_18054502 |
| A/dolphin/Peru/PIU-002/2022 | EPI_ISL_18054503 |
| A/sea lion/Peru/AQP-SER00B/2023 | EPI_ISL_18054508 |
| A/sea lion/Peru/AQP-SER00K/2023 | EPI_ISL_18054509 |
| A/sea lion/Peru/AQP-SER00R/2023 | EPI_ISL_18054510 |
| A/Thalasseus_maximus/Brazil-ES/23ES1A0008/2023 | EPI_ISL_18130597 |
| A/Thalasseus_acuflavidus/Brazil-ES/23ES1A0009/2023 | EPI_ISL_18130622 |
| A/Thalasseus_acuflavidus/Brazil-ES/23ES1A0025/2023 | EPI_ISL_18130627 |
| A/Thalasseus_maximus/Brazil-ES/23ES1A0026/2023 | EPI_ISL_18130628 |
| A/American Wigeon/South Carolina/22-000345-001/2021 | EPI_ISL_18133029 |
| A/Common Raven/Connecticut/USDA-009204-001/2022 | EPI_ISL_18133350 |
| A/wildbird-Sula-nebouxii/Ecuador/7607/2023 | EPI_ISL_18137626 |
| A/duck/Peru/CAL-INS-013/2023 | EPI_ISL_18217104 |
| A/american kestrel/Peru/UNMSM-A273/2022 | EPI_ISL_18238548 |
| A/peruvian pelican/Peru/UNMSM-A106/2022 | EPI_ISL_18238607 |
| A/belchers gull/Peru/UNMSM-A267/2022 | EPI_ISL_18238609 |
| A/south american sea lion/Peru/AQP-SER00K/2023 | EPI_ISL_18265321 |
| A/dolphin/Peru/PIU-SER002/2022 | EPI_ISL_18265431 |
| A/Sanderling/Peru/PIU-SER005/2022 | EPI_ISL_18265432 |
| A/pelican/Peru/PIU-SER013/2022 | EPI_ISL_18265433 |
| A/Guanay cormorant/Peru/PIU-SER024/2022 | EPI_ISL_18265434 |
| A/pelican/Peru/PIU-SER019/2022 | EPI_ISL_18265435 |
| A/pelican/Peru/PIU-SER028/2022 | EPI_ISL_18265436 |
| A/pelican/Peru/PIU-SER016/2022 | EPI_ISL_18265437 |

| | |
|---|---|
| A/backyard chicken/Uruguay/UDELAR-040-M5/2023 | EPI_ISL_18310942 |
| A/black-necked swan/Uruguay/UDELAR-078-M2/2023 | EPI_ISL_18310957 |
| A/black-necked swan/Uruguay/UDELAR-014-M3/2023 | EPI_ISL_18310958 |
| A/backyard turkey/Uruguay/UDELAR-124-M6/2023 | EPI_ISL_18310959 |
| A/backyard duck/Uruguay/UDELAR-124-M3/2023 | EPI_ISL_18310960 |
| A/backyard chicken/Uruguay/UDELAR-144-M3/2023 | EPI_ISL_18310961 |
| A/backyard chicken/Uruguay/UDELAR-127-M4/2023 | EPI_ISL_18310962 |
| A/backyard chicken/Uruguay/UDELAR-127-M1/2023 | EPI_ISL_18310963 |
| A/backyard chicken/Uruguay/UDELAR-124-M1/2023 | EPI_ISL_18310964 |
| A/backyard chicken/Uruguay/UDELAR-047-M3/2023 | EPI_ISL_18310965 |
| A/backyard chicken/Uruguay/UDELAR-047-M1/2023 | EPI_ISL_18310966 |
| A/backyard chicken/Uruguay/UDELAR-040-M7/2023 | EPI_ISL_18310967 |
| A/Belcher gull/Peru/UNMSM-A267/2022 | EPI_ISL_18371664 |
| A/Guanay cormorant/Peru/UNMSM-A275/2022 | EPI_ISL_18371665 |
| A/Peruvian booby/Peru/UNMSM-A296/2022 | EPI_ISL_18371666 |
| A/Brown_skua/Bird_Island/128287/2023 | EPI_ISL_18439562 |
| A/Brown_skua/Bird_Island/128288/2023 | EPI_ISL_18439563 |
| A/Brown_skua/Bird_Island/128289/2023 | EPI_ISL_18439564 |
| A/duck/Peru/LAM-INS-014/2023 | EPI_ISL_18497946 |
| A/Southern_fulmar/Falkland_Islands/133789/2023 | EPI_ISL_18522961 |
| A/Kelp_Gull/Hound_Bay/133744/2023 | EPI_ISL_18592422 |
| A/Kelp_Gull/Hound_Bay/133747/2023 | EPI_ISL_18592423 |
| A/Brown_Skua/Moltke_Harbour/133752/2023 | EPI_ISL_18592424 |
| A/Kelp_Gull/Moltke_Harbour/133754/2023 | EPI_ISL_18592425 |
| A/Brown_Skua/Moltke_Harbour/133755/2023 | EPI_ISL_18592426 |
| A/Kelp_Gull/Harpon_Bay/133943/2023 | EPI_ISL_18592427 |
| A/Brown_Skua/Hound_Bay/133947/2023 | EPI_ISL_18592428 |
| A/Brown_Skua/Hound_Bay/133949/2023 | EPI_ISL_18592429 |
| A/Band-tailed gull/Tarapaca/236339/2023 | EPI_ISL_18690676 |
| A/chicken/Maule/242133-1/2023 | EPI_ISL_18690677 |
| A/chicken/Metropolitana/245704-1/2023 | EPI_ISL_18690679 |
| A/chicken/Nuble/241585/2023 | EPI_ISL_18690680 |
| A/chicken/OHiggins/242581-2/2023 | EPI_ISL_18690682 |
| A/cormorant/Antofagasta/236025-1/2023 | EPI_ISL_18690683 |
| A/domestic duck/Araucania/SJCEIRR-2439541/2023 | EPI_ISL_18690684 |
| A/goose/Araucania/244373-1 /2023 | EPI_ISL_18690687 |
| A/kelp gull/Nuble/234619-1/2023 | EPI_ISL_18690704 |
| A/turkey vulture/Atacama/241294/2023 | EPI_ISL_18690710 |
| A/chicken/Bio Bio/245039-2/2023 | EPI_ISL_18690712 |
| A/chicken/Bio Bio/241781/2023 | EPI_ISL_18690713 |
| A/Peruvian booby/Antofagasta/236109-1/2023 | EPI_ISL_18690723 |

| | |
|---|---|
| A/Peruvian booby/Arica y Parinacota/234906-1/2023 | EPI_ISL_18690725 |
| A/Peruvian booby/Tarapaca/236308/2023 | EPI_ISL_18690730 |
| A/chicken/Araucania/SJCEIRR-245202/2023 | EPI_ISL_18690740 |
| A/Peruvian booby/Valparaiso/236604/2023 | EPI_ISL_18690744 |
| A/Goose/Argentina/389-1/2023 | EPI_ISL_18698459 |
| A/Chicken/Argentina/464-4/2023 | EPI_ISL_18698460 |
| A/Chicken/Argentina/477-5/2023 | EPI_ISL_18698462 |
| A/Chicken/Argentina/481-2/2023 | EPI_ISL_18698464 |
| A/Chicken/Argentina/485-6/2023 | EPI_ISL_18698466 |
| A/Chicken/Argentina/491-2/2023 | EPI_ISL_18698469 |
| A/Chicken/Argentina/501-1/2023 | EPI_ISL_18698471 |
| A/Chicken/Argentina/506-2/2023 | EPI_ISL_18698474 |
| A/Chicken/Argentina/509-2/2023 | EPI_ISL_18698476 |
| A/Chicken/Argentina/556-6/2023 | EPI_ISL_18698478 |
| A/Chicken/Argentina/559-8/2023 | EPI_ISL_18698480 |
| A/Chicken/Argentina/578-2/2023 | EPI_ISL_18698482 |
| A/Avian/Argentina/579-5/2023 | EPI_ISL_18698485 |
| A/Avian/Argentina/586-4/2023 | EPI_ISL_18698487 |
| A/Chicken/Argentina/588-4/2023 | EPI_ISL_18698490 |
| A/Chicken/Argentina/606-1/2023 | EPI_ISL_18698492 |
| A/Chicken/Argentina/736-1/2023 | EPI_ISL_18698494 |
| A/Turkey/Argentina/753-1/2023 | EPI_ISL_18698497 |
| A/Chicken/Argentina/858-1/2023 | EPI_ISL_18698499 |
| A/Chicken/Argentina/919-3/2023 | EPI_ISL_18698502 |
| A/Chicken/Argentina/1035-1/2023 | EPI_ISL_18698504 |
| A/Chicken/Argentina/1147-2/2023 | EPI_ISL_18698505 |
| A/Chicken/Argentina/1200-1/2023 | EPI_ISL_18698506 |
| A/Chicken/Argentina/1340-2/2023 | EPI_ISL_18698507 |
| A/Turkey/Argentina/1348-3/2023 | EPI_ISL_18698508 |
| A/Chicken/Argentina/1375-6/2023 | EPI_ISL_18698509 |
| A/Chicken/Argentina/1416-3/2023 | EPI_ISL_18698510 |
| A/Chicken/Argentina/1530-3/2023 | EPI_ISL_18698511 |
| A/Chicken/Argentina/1708-1/2023 | EPI_ISL_18698512 |
| A/Turkey/Argentina/1710-1/2023 | EPI_ISL_18698513 |
| A/Turkey/Argentina/1711-2/2023 | EPI_ISL_18698514 |
| A/Duck/Argentina/1712-5/2023 | EPI_ISL_18698515 |
| A/Avian/Argentina/1762-2/2023 | EPI_ISL_18698516 |
| A/Avian/Argentina/1790-5/2023 | EPI_ISL_18698517 |
| A/Chicken/Argentina/1976-2/2023 | EPI_ISL_18698518 |
| A/Chicken/Argentina/1984-4/2023 | EPI_ISL_18698519 |
| A/Chicken/Argentina/2034-5/2023 | EPI_ISL_18698520 |

| | |
|---|---|
| A/Chicken/Argentina/2049-3/2023 | EPI_ISL_18698521 |
| A/Chicken/Argentina/2064-3/2023 | EPI_ISL_18698522 |
| A/Duck/Argentina/2197-1/2023 | EPI_ISL_18698523 |
| A/Chicken/Argentina/2305-1/2023 | EPI_ISL_18698524 |
| A/Chicken/Argentina/2483-3/2023 | EPI_ISL_18698526 |
| A/Chicken/Argentina/2796-2/2023 | EPI_ISL_18698527 |
| A/Chicken/Argentina/3346-1/2023 | EPI_ISL_18698528 |
| A/Chicken/Argentina/3657-2/2023 | EPI_ISL_18698529 |
| A/Chicken/Argentina/3695-3/2023 | EPI_ISL_18698530 |
| A/Chicken/Argentina/1495-4/2023 | EPI_ISL_18698719 |
| A/Chicken/Argentina/747-1/2023 | EPI_ISL_18698730 |
| A/Chicken/Argentina/895-1/2023 | EPI_ISL_18698731 |
| A/Chicken/Argentina/2016-2/2023 | EPI_ISL_18698732 |
| A/Sea Lion/Argentina/3849-4/2023 | EPI_ISL_18698754 |
| A/Sea Lion/Argentina/3893-1/2023 | EPI_ISL_18698755 |
| A/goose/France/PPNL-21P007579/2021 | EPI_ISL_18718155 |
| A/Antarctic_Fur_Seal/Jason_Harbour/141037/2023 | EPI_ISL_18742212 |
| A/Southern_Elephant_Seal/Jason_Harbour/141078/2023 | EPI_ISL_18742213 |
| A/Brown_Skua/Bird_Island/141232/2023 | EPI_ISL_18742214 |
| A/Kelp_Gull/Penguin_River/141234/2023 | EPI_ISL_18742215 |
| A/Brown_Skua/Penguin_River/141236/2023 | EPI_ISL_18742216 |
| A/Kelp_Gull/Penguin_River/141239/2023 | EPI_ISL_18742217 |
| A/Brown_Skua/Penguin_River/141240/2023 | EPI_ISL_18742218 |
| A/South_Georgia_Shag/King_Edward_Cove/141245/2023 | EPI_ISL_18742219 |
| A/Antarctic_tern/King_Edward_Point/141271/2023 | EPI_ISL_18742220 |
| A/Southern_Elephant_Seal/Jason_Harbour/141027/2023 | EPI_ISL_18742221 |
| A/frigatebird/Rio de Janeiro/MAPA-1532N/2023 | EPI_ISL_18755251 |
| A/tern/Espirito Santo/MAPA-1339N2/2023 | EPI_ISL_18755338 |
| A/chicken/Metropolitana/243148-1/2023 | EPI_ISL_18760060 |
| A/pelican/Bio Bio/236574/2023 | EPI_ISL_18760061 |
| A/turkey/Valparaiso/245562-1/2023 | EPI_ISL_18760063 |
| A/chicken/Aysen/SJCEIRR-2477921/2023 | EPI_ISL_18760064 |
| A/caracara/Coquimbo/SJCEIRR-2423261/2023 | EPI_ISL_18760065 |
| A/Humboldt penguin/Antofagasta/236063-2/2023 | EPI_ISL_18760066 |
| A/Peruvian booby/OHiggins/242755-1/2023 | EPI_ISL_18760067 |
| A/chicken/Aysen/SJCEIRR-2477922/2023 | EPI_ISL_18760068 |
| A/sea lion/Valparaiso/SJCEIRR-2431361/2023 | EPI_ISL_18760069 |
| A/black-necked swan/Los Rios/247292-1/2023 | EPI_ISL_18760070 |
| A/brown-hooded gull/Los Rios/247093-1/2023 | EPI_ISL_18760071 |
| A/brown-hooded gull/Los Rios/247094-1/2023 | EPI_ISL_18760072 |
| A/chicken/Araucania/244469/2023 | EPI_ISL_18760073 |

| | |
|---|---|
| A/chicken/Atacama/235254-2/2023 | EPI_ISL_18760074 |
| A/sea lion/Brazil/KU-3581/2023 | EPI_ISL_18773600 |
| A/sea lion/Brazil/KU-3591/2023 | EPI_ISL_18773668 |
| A/porpoise/Antofagasta/SJCEIRR-2465061/2023 | EPI_ISL_18777129 |
| A/gull/Maule/234329-1/2023 | EPI_ISL_18777130 |
| A/dolphin/Nuble/SJCEIRR-2482441/2023 | EPI_ISL_18777138 |
| A/dolphin/Maule/SJCEIRR-246026/2023 | EPI_ISL_18777139 |
| A/porpoise/Atacama/SJCEIRR-245355/2023 | EPI_ISL_18777140 |
| A/porpoise/Antofagasta/SJCEIRR-2465062/2023 | EPI_ISL_18777141 |
| A/Pelecanus occidentalis/Venezuela/Inf1/2022 | EPI_ISL_18909019 |
| A/Southern_Fulmar/Falkland_Islands/138870/2023 | EPI_ISL_18918640 |
| A/Pelecanus occidentalis/Venezuela/Inf3/2022 | EPI_ISL_18931239 |
| A/Pelecanus occidentalis/Venezuela/Inf10/2022 | EPI_ISL_18931240 |
| A/Pelecanus occidentalis/Venezuela/Inf11/2022 | EPI_ISL_18931241 |
| A/Pelecanus occidentalis/Venezuela/Inf14/2022 | EPI_ISL_18931268 |
| A/Pelecanus occidentalis/Venezuela/Inf15/2022 | EPI_ISL_18931269 |
| A/Pelecanus occidentalis/Venezuela/Inf17/2022 | EPI_ISL_18931270 |
| A/Pelecanus occidentalis/Venezuela/Inf19/2022 | EPI_ISL_18931271 |
| A/Pelecanus occidentalis/Venezuela/Inf20/2022 | EPI_ISL_18931272 |
| A/Coragyps atratus/Venezuela/Inf21/2022 | EPI_ISL_18931273 |
| A/Pelecanus occidentalis/Venezuela/Inf41/2023 | EPI_ISL_18931274 |
| A/Pelecanus occidentalis/Venezuela/Inf43/2023 | EPI_ISL_18931275 |
| A/Pelecanus occidentalis/Venezuela/Inf44/2023 | EPI_ISL_18931276 |
| A/Franklin's gull/OHiggins/236195-1/2023 | EPI_ISL_18939722 |
| A/South American tern/Argentina/RN-PB015/2023 | EPI_ISL_18945315 |
| A/South American sea lion/Argentina/RN-PB013/2023 | EPI_ISL_18945316 |
| A/South American sea lion/Argentina/RN-PB011/2023 | EPI_ISL_18945317 |
| A/South American sea lion/Argentina/RN-PB007/2023 | EPI_ISL_18945319 |
| A/South American fur seal/Argentina/RN-PB019/2023 | EPI_ISL_18945320 |
| A/Sula_leucogaster/Brazil-PR/A0001-IBTEC/2023 | EPI_ISL_18975372 |
| A/Est/Parnu/eagle/TA21-11864-1/2021 | EPI_ISL_19028663 |
| A/Calidris alba/Lima/Villa01/2023 (this) | EPI_ISL_19070288 |
| A/pinniped/Uruguay/P10_8923/2023 | EPI_ISL_19070481 |
| A/pinniped/Uruguay/P13_11923/2023 | EPI_ISL_19070482 |
| A/pinniped/Uruguay/P14_11923/2023 | EPI_ISL_19070483 |
| A/pinniped/Uruguay/P15_14923/2023 | EPI_ISL_19070484 |
| A/pinniped/Uruguay/P17_14923/2023 | EPI_ISL_19070485 |
| A/pinniped/Uruguay/P18_14923/2023 | EPI_ISL_19070486 |
| A/pinniped/Uruguay/P26_21023/2023 | EPI_ISL_19070488 |
| A/pinniped/Uruguay/P7_6923/2023 | EPI_ISL_19070491 |
| A/pinniped/Uruguay/P8_8923/2023 | EPI_ISL_19070492 |

| | |
|---|---|
| A/seabird/Uruguay/P16_14923/2023 | EPI_ISL_19070493 |
| A/seabird/Uruguay/P23_41023/2023 | EPI_ISL_19070495 |
| A/seabird/Uruguay/P24_41023/2023 | EPI_ISL_19070496 |
| A/seabird/Uruguay/P25_41023/2023 | EPI_ISL_19070497 |
| A/pinniped/Uruguay/P4_6923/2023 | EPI_ISL_19070498 |
| A/Inca tern/Antofagasta/238083/2023 | EPI_ISL_19131156 |
| A/Inca tern/Arica y Parinacota/227519-1/2022 | EPI_ISL_19131159 |
| A/Inca tern/Coquimbo/239619/2023 | EPI_ISL_19131162 |
| A/Peruvean booby/Valparaiso/238535/2023 | EPI_ISL_19131163 |
| A/Peruvian booby/Atacama/230579-1/2023 | EPI_ISL_19131164 |
| A/Peruvian booby/OHiggins/234887-1/2023 | EPI_ISL_19131165 |
| A/Band-tailed gull/Antofagasta/228525-1/2022 | EPI_ISL_19131166 |
| A/Band-tailed gull/Arica y Parinacota/232200-1/2023 | EPI_ISL_19131168 |
| A/band-tailed gull/Tarapaca/238807-1/2023 | EPI_ISL_19131176 |
| A/great grebe/Atacama/231482-1/2023 | EPI_ISL_19131195 |
| A/great grebe/Atacama/231482-2/2023 | EPI_ISL_19131196 |
| A/Peruvean booby/Atacama/230579-1/2023 | EPI_ISL_19139974 |
| A/guanay cormorant/Atacama/235520-1/2023 | EPI_ISL_19151400 |
| A/Franklin gull/OHiggins/236195-1/2023 | EPI_ISL_19162729 |
| A/Great grabe/Atacama/231482-1/2023 | EPI_ISL_19162730 |
| A/Great grabe/Atacama/231482-2/2023 | EPI_ISL_19162731 |
| A/Buteogallus urubitinga/Espirito Santo/1630-SC/2023 | EPI_ISL_19215175 |
| A/Gallus gallus/Mato Grosso do Sul/2108-SN52/2023 | EPI_ISL_19215176 |
| A/Megascops choliba/Espirito Santo/1409-N/2023 | EPI_ISL_19215177 |
| A/Numida meleagris/Santa Catarina/1843-N3/2023 | EPI_ISL_19215178 |
| A/Otaria flavescens/Rio Grande do Sul/2148-N/2023 | EPI_ISL_19215179 |
| A/Otaria flavescens/Rio Grande do Sul/2165-SO/2023 | EPI_ISL_19215180 |
| A/Pluvialis dominica/Sao Paulo/2252-N/2023 | EPI_ISL_19215181 |
| A/Procellaria aequinoctialis/Sao Paulo/2259-N/2023 | EPI_ISL_19215182 |
| A/Sterna hirundo/Espirito Santo/1455-N/2023 | EPI_ISL_19215183 |
| A/Sterna hirundo/Santa Catarina/2261-N/2023 | EPI_ISL_19215184 |
| A/Sula leucogaster/Sao Francisco do Sul/2122-N/2023 | EPI_ISL_19215185 |
| A/Thalasseus maximus/Parana/1775-N/2023 | EPI_ISL_19215186 |
| A/Thalasseus maximus/Santa Catarina/1941-N/2023 | EPI_ISL_19215187 |
| A/Thalasseus maximus/Rio Grande do Sul/2177-N/2023 | EPI_ISL_19215188 |
| A/Thalasseus maximus/Sao Paulo/1546-N/2023 | EPI_ISL_19215189 |
| A/Procellaria aequinoctialis/Sao Paulo/2271-N/2023 | EPI_ISL_19215190 |
| A/Sterna hirundo/Rio de Janeiro/0613-N/2024 | EPI_ISL_19215191 |
| A/Sterna hirundo/Espirito Santo/0155-N/2024 | EPI_ISL_19215192 |
| A/Sterna hirundo/Rio de Janeiro/0177-N/2024 | EPI_ISL_19215193 |
| A/Sterna hirundo/Espirito Santo/0448-N/2024 | EPI_ISL_19215194 |

| | |
|---|---|
| A/Sterna hirundo/Rio de Janeiro/0721-N/2024 | EPI_ISL_19215195 |
| A/Sterna hirundo/Rio de Janeiro/0481-R/2024 | EPI_ISL_19215196 |
| A/Thalasseus acuflavidus/Sao Paulo/0291-N/2024 | EPI_ISL_19215197 |
| A/Thalasseus acuflavidus/Sao Paulo/2280-N/2023 | EPI_ISL_19215198 |
| A/Thalasseus acuflavidus/Parana/2277-N/2023 | EPI_ISL_19215199 |
| A/Thalasseus maximus/Sao Paulo/2339-N/2023 | EPI_ISL_19215200 |
| A/burmeisters porpoise/Chile/246506-2/2023 | EPI_ISL_19391458 |
| A/chungungo/Chile/241436/2023 | EPI_ISL_19391459 |
| A/Guanay cormorant/Chile/239584/2023 | EPI_ISL_19391460 |
| A/pelican/Chile/229424-4/2022 | EPI_ISL_19391461 |
| A/sea lion/Atacama/242444-1/2023 | EPI_ISL_19391462 |
| A/Pelican/Coquimbo/233565-1/2023 | EPI_ISL_19404893 |
| A/turkey/Valparaiso/248976-12/2023 | EPI_ISL_19404902 |
| A/kelp gull/Valparaiso/236011-1/2023 | EPI_ISL_19404906 |
| A/gray gull/Arica y Parinacota/235908-1/2023 | EPI_ISL_19404913 |
| A/chicken/Aysen/250755-2/2023 | EPI_ISL_19404916 |
| A/sea lion/Los Lagos/247932/2023 | EPI_ISL_19404920 |
| A/Sanderling/Chile/230758-2/2023 | EPI_ISL_19404931 |
| A/turkey/Valparaiso/248976-13/2023 | EPI_ISL_19404939 |
| A/chicken/Cochabamba/39356/2023 | EPI_ISL_19410267 |
| A/chicken/Cochabamba/39383/2023 | EPI_ISL_19410268 |
| A/chicken/Cochabamba/39425/2023 | EPI_ISL_19410269 |
| A/chicken/Cochabamba/39472/2023 | EPI_ISL_19410270 |
| A/chicken/Cochabamba/39476/2023 | EPI_ISL_19410271 |
| A/chicken/Cochabamba/39507/2023 | EPI_ISL_19410272 |
| A/chicken/Cochabamba/39522/2023 | EPI_ISL_19410273 |
| A/Chilean teal/Nuble/233516-2/2023 | EPI_ISL_19410275 |
| A/peruvian booby/Coquimbo/242304/2023 | EPI_ISL_19410277 |
| A/Sanderling/Arica y Parinacota/230758-2/2022 | EPI_ISL_19410278 |
| A/sea lion/Tarapaca/245719-10/2023 | EPI_ISL_19410279 |
| A/chicken/Maule/247588-1/2023 | EPI_ISL_19410280 |
| A/gray gull/Arica/239053-1/2023 | EPI_ISL_19410281 |
| A/kelp gull/Atacama/234251-1/2023 | EPI_ISL_19410282 |
| A/peregrine falcon/Antofagasta/235144-1/2023 | EPI_ISL_19410283 |
| A/turkey/Valparaiso/245886-1/2023 | EPI_ISL_19410284 |
| A/turkey/Valparaiso/245886-3/2023 | EPI_ISL_19410285 |
| A/turkey/Valparaiso/245886-5/2023 | EPI_ISL_19410286 |
| A/chicken/Cochabamba/39391/2023 | EPI_ISL_19410294 |
| A/chicken/Cochabamba/39404/2023 | EPI_ISL_19410295 |
| A/chicken/Cochabamba/39521/2023 | EPI_ISL_19410296 |
| A/chicken/Cochabamba/39555/2023 | EPI_ISL_19410297 |

| | |
|---|---|
| A/chicken/Cochabamba/39581/2023 | EPI_ISL_19410298 |
| A/chicken/Potosi/39870/2023 | EPI_ISL_19410299 |
| A/gull/Biobio/237012/2023 | EPI_ISL_19410300 |
| A/marine otter/Valparaiso/SP010/2023 | EPI_ISL_19418468 |
| A/chicken/Cochabamba/39539/2023 | EPI_ISL_19418487 |
| A/river otter/Magallanes/257113-1/2023 | EPI_ISL_19418489 |
| A/Chilean skua/Magallanes/248128-1/2023 | EPI_ISL_19418490 |
| A/buff-necked ibis/Bio bio/247636-1/2023 | EPI_ISL_19418491 |
| A/slender-billed parakeet/Araucania/242881-1/2023 | EPI_ISL_19422674 |
| A/South American tern/Argentina/CH-PD030/2023 | EPI_ISL_19466158 |
| A/Southern elephant seal/Peninsula Valdes/PB2_CH-PD027/2023 | EPI_ISL_19466175 |
| A/royal tern/Argentina/CH-PD036/2023 | EPI_ISL_19466181 |
| A/southern elephant seal/Argentina/CH-PD035/2023 | EPI_ISL_19466183 |
| A/South American tern/Argentina/CH-PD037/2023 | EPI_ISL_19466210 |
| A/Southern elephant seal/Peninsula Valdes/NA_CH-PD032_lung/2023 | EPI_ISL_19466224 |
| A/Southern elephant seal/Peninsula Valdes/NA_CH-PD032_oral/2023 | EPI_ISL_19466225 |
| A/Southern elephant seal/Peninsula Valdes/NA_CH-PD032_tracheal/2023 | EPI_ISL_19466227 |
| A/Southern elephant seal/Peninsula Valdes/NA_CH-PD032rain/2023 | EPI_ISL_19466228 |
| A/southern elephant seal/Argentina/CH-PM053/2023 | EPI_ISL_19466252 |
| A/California/194/2024 | EPI_ISL_19616453 |
| A/Fujian-Sanyuan/21099/2017 | EPI_ISL_8767096 |
| A/chicken/Nigeria/VRD21-102_21VIR2370-424/2021_HA | MW961452.1 |
| A/chicken/Nigeria/VRD21-109_21VIR2370-425/2021_HA | MW961460.1 |
| A/chicken/Lesotho/341 | OL477469.1 |
| A/snow goose/Kansas/W22-199D/2022_HA | OP221285.1 |
| A/blue-winged teal/Texas/UGAI22-3189/2022_PB2 | OQ733035.1 |
| A/Peruvian_pelican/Peru/A074/2022_HA | OQ747758.1 |
| A/Belcher's_gull/Peru/A102/2022_HA | OQ747759.1 |
| A/Belcher's_gull/Peru/A267/2022_HA | OQ747761.1 |
| A/Whimbrel/Coquimbo/239964/2023_HA | OR125135.1 |
| A/Pelican/Maule/231155-2/2023_HA | OR125307.1 |
| A/Turkey vulture/Valparaiso/230187-1/2022_HA | OR125345.1 |
| A/South American sea lion/Tarapaca/240524-2/2023_HA | OR125370.1 |
| A/Turkey vulture/Antofagasta/228252-1/2022_HA | OR265539.1 |
| A/Otaria flavescens/Torres/2165-SO/2023_HA | OR852415.1 |
| A/gray gull/Atacama/235521-1/2023_PB2 | OR910143.1 |
| A/cormorant/Antofagasta/236025-1/2023_HA | OR910159.1 |
| A/Peruvian pelican/Nuble/236068-1/2023_PB2 | OR910176.1 |
| A/Band-tailed gull/Tarapaca/236339/2023_HA | OR910186.1 |
| A/pelican/Arica y Parinacota/235876-4/2023_PB2 | OR910221.1 |
| A/pelican/Arica y Parinacota/235876-4/2023_NA | OR910225.1 |

| | |
|---|---|
| A/chicken/Atacama/235254-6/2023_PB2 | OR910228.1 |
| A/Chimango caracara/Coquimbo/242326-1/2023_PB2 | OR910242.1 |
| A/Chimango caracara/Coquimbo/242326-1/2023_HA | OR910243.1 |
| A/Turkey vulture/Antofagasta/236643/2023_PB2 | OR910259.1 |
| A/Turkey vulture/Antofagasta/236643/2023_HA | OR910261.1 |
| A/chicken/Maule/242947-1/2023_NA | OR910283.1 |
| A/Peruvian booby/Coquimbo/238918/2023_PB2 | OR910300.1 |
| A/Peruvian booby/Coquimbo/238918/2023_HA | OR910301.1 |
| A/Peruvian booby/Coquimbo/238918/2023_NA | OR910303.1 |
| A/Peruvian booby/Antofagasta/236109-1/2023_NA | OR910313.1 |
| A/band-tailed gull/Arica y Parinacota/235901-2/2023_HA | OR910318.1 |
| A/Peruvian booby/Coquimbo/239024/2023_NA | OR910368.1 |
| A/gray gull/Antofagasta/236047-1/2023_PB2 | OR910369.1 |
| A/gray gull/Antofagasta/236047-1/2023_HA | OR910371.1 |
| A/gray gull/Antofagasta/236047-1/2023_NA | OR910373.1 |
| A/chicken/Atacama/235254-5/2023_NA | OR910398.1 |
| A/Peruvian booby/Tarapaca/236308/2023_HA | OR910404.1 |
| A/chicken/Bio Bio/245039-2/2023_NA | OR910437.1 |
| A/gray gull/Tarapaca/236305/2023_PB2 | OR910446.1 |
| A/gray gull/Tarapaca/236305/2023_HA | OR910447.1 |
| A/South American sea lion/Valparaiso/243136-1/2023_HA | OR910468.1 |
| A/South American sea lion/Valparaiso/244738-1/2023_HA | OR910481.1 |
| A/Peruvian booby/Arica y Parinacota/234906-1/2023_PB2 | OR910507.1 |
| A/penguin/Antofagasta/234905-1/2023_NA | OR910517.1 |
| A/seabird/Uruguay/P16_14926/2023_HA | OR912290.1 |
| A/seabird/Uruguay/P25_41023/2023_HA | OR912294.1 |
| A/pinniped/Uruguay/P26_21023/2023_HA | OR912295.1 |
| A/pinniped/Uruguay/P7_6923/2023_NA | OR912312.1 |
| A/seabird/Uruguay/P25_41023/2023_NA | OR912321.1 |
| A/Chilean dolphin/Nuble/248244-1/2023_HA | OR960980.1 |
| A/guanay cormorant/Tarapaca/236301/2023_PB2 | OR960990.1 |
| A/guanay cormorant/Tarapaca/236301/2023_HA | OR960992.1 |
| A/guanay cormorant/Atacama/235246-1/2023_HA | OR960998.1 |
| A/guanay cormorant/Arica y Parinacota/235902-2/2023_PB2 | OR961003.1 |
| A/guanay cormorant/Arica y Parinacota/235902-2/2023_HA | OR961005.1 |
| A/guanay cormorant/Coquimbo/239080/2023_PB2 | OR961009.1 |
| A/South American sea lion/Bio Bio/246296-1/2023_HA | OR979626.1 |
| A/South American sea lion/Argentina/RN-PB004/2023_PB2 | OR987081.1 |
| A/South American sea lion/Argentina/RN-PB007/2023_HA | OR987092.1 |
| A/South American sea lion/Brazil/OF-358_4/2023_HA | PP094671.1 |
| A/South American sea lion/Brazil/OF-359_4/2023_HA | PP094679.1 |

| | |
|---|---|
| A/Peruvian booby/OHiggins/234887-1/2023_HA | PP757802.1 |
| A/band-tailed gull/Antofagasta/228525-2/2022_HA | PP758168.1 |
| A/Procellaria aequinoctialis/UbatubaBR/2271-N/2023_HA | PP786400.1 |
| A/Southern Fulmar/South Georgia and the South Sandwich Islands/9/2023_PB2 | PQ113938.1 |
| A/Southern Fulmar/South Georgia and the South Sandwich Islands/12/2023_HA | PQ113941.1 |
| A/Sanderling/CHL/230758-2/2023_HA | PQ304209.1 |

**Supplementary Table 3.** GISAID acknowledgment table under GISAID Identifier: EPI_SET_241228nx and DOI: 10.55876/gis8.241228nx. All genome sequences and associated metadata in this dataset are published in GISAID's EpiFlu database. To view the contributors for each individual sequence, along with details such as accession number, virus name, collection date, originating lab, submitting lab, and list of authors, visit DOI: 10.55876/gis8.241228nx. EPI_SET_241228nx comprises 425 individual viruses, with collection dates ranging from 2017-01-01 to 2024-11-22. Data were collected from 20 countries and territories.
